# The contribution of stimulating multiple body parts simultaneously to the illusion of owning an entire artificial body

**DOI:** 10.1101/2020.05.04.076497

**Authors:** Sophie H. O’Kane, H. Henrik Ehrsson

**Affiliations:** Department of Neuroscience, Karolinska Institutet, Stockholm, Sweden

## Abstract

The full-body ownership illusion exploits multisensory perception to induce a feeling of ownership for an entire artificial body. Whilst previous research has shown that the synchronous visuotactile stimulation of a single body part is sufficient for illusory ownership over the whole body, the effect of combining multisensory stimulation across multiple body parts remains unknown. Therefore, 48 healthy adults participated in conditions of a full-body ownership illusion involving synchronous or asynchronous visuotactile stimulation to one, two or three body parts simultaneously (2 x 3 design). We developed a novel questionnaire to isolate the sense of ownership of five specific body parts (left leg, right leg, left arm, right arm, and trunk) from the full-body ownership experience and sought not only to test for greater (part and whole) body ownership in synchronous versus asynchronous stimulation, but also, potentially varying degrees of illusion intensity related to the number of body parts stimulated. As expected, illusory full-body ownership and all five body-part ownership ratings were significantly higher following synchronous stimulation (all p values ≤.01). Since non-stimulated body parts also received significantly higher ownership ratings following synchronous stimulation, the results are consistent with an illusion engaging the entire body. We further noted that ownership ratings for the right body parts (often stimulated) were significantly higher than ownership ratings for the left body parts (never stimulated). Regarding explicit feelings of full-body ownership, subjective ratings were not significantly enhanced by increasing the number of synchronously stimulated body parts (synchronicity x number stimulated interaction; p.099). Instead, median ratings indicated a moderate affirmation (+1) of full-body illusory sensation for all three synchronous conditions; a finding mirrored by full-body illusion onset time. The results support the notion that feelings of full-body ownership are mediated by a generalisation from stimulated part(s)-to-whole, supported by processes related to multisensory body perception.

## Introduction

How do we come to perceive the body, the single, integrated biological entity in which we sense and act upon our world, as belonging exclusively to oneself? What are the neurocognitive principles governing the perception of our own body not as a set of fragmented segments, but as the gestalt that delineates the boundaries between what is me versus what is not? The feeling of ‘body ownership’ (Makin et al. 2008; Ehrsson 2012, 2020; Blanke et al. 2015; Gallagher, 2000; Tsakiris 2010) attracts attention across diverse academic fields, although the distinction between part and whole in body ownership, herein referred to as ‘body-part ownership’ and ‘full-body ownership’, respectively (Ehrsson, 2012; 2020; Gentile et al., 2015; Petkova, Björnsdotter, et al., 2011; Petkova & Ehrsson, 2008), has been studied less often. In cognitive neuroscience, the discovery of the rubber hand Illusion (Botvinick & Cohen, 1998) led to an exciting expanse in empirical research toward understanding the perceptual processes and neural mechanisms that underpin ownership of a single limb in healthy individuals. Using experimental conditions to exploit the basic principles of multisensory integration (Stein & Stanford, 2008), this simple perceptual illusion provides an exquisite demonstration of the malleability of the sense of body ownership amongst healthy people (Ehrsson, 2012, 2020). However, in addition to inducing a sense of ownership for a prosthetic hand, Petkova and Ehrsson (2008) revealed that the illusory experience of ownership could also be extended to encompass an entire artificial body, which opened up the avenue for conducting experimental research on full-body ownership alongside body-part ownership.

During the ‘full-body ownership illusion’ (also referred to as the ‘body-swap illusion’, Petkova & Ehrsson, 2008), tactile stimulation is administered to the participant’s real body, in perfect spatio-temporal synchrony with visual feedback of identical stimuli being applied to a fake plastic mannequin’s body. For this particular paradigm, the artificial body must be presented in an anatomically congruent position from the natural first-person point of view, or as mirror reflection, easily achieved using modern head mounted displays connected to video cameras (Petkova, Khoshnevis & Ehrsson, 2011; Preston, Kuper-Smith & Ehrsson, 2015). Subjective reports of referral of touch, illusory experiences, so as to directly feel the touches applied to the artificial body, plus some degree of illusory ownership over its entirety, are both well-supported in the majority of participants (Petkova & Ehrsson, 2008; Slater, Perez-Marcos, Ehrsson & Sanchez-Vives, 2009; Petkova et al., 2011; Gentile, Björnsdotter, Petkova, Abdulkarim & Ehrsson, 2015). These results support multisensory integration, namely that of visual, tactile and proprioceptive input, as an essential framework to investigate the feeling of full-body ownership (Ehrsson, 2012, 2020; Tsakiris, 2017; Kilteni et al. 2015).

In order for the illusory percept of ownership to arise, multisensory stimulation must obey basic rules for successful integration. Visuotactile stimuli must be temporally synchronous, whilst asynchronous visuotactile stimulation provides a reliable control condition to the majority of studies (Petkova & Ehrsson, 2008; Petkova et al. 2011; Gentile et al., 2015; Guterstam et al. 2015; Preston et al., 2015; Kokkinara & Slater, 2014). Moreover, visuotactile stimulation must be spatially congruent, i.e. applied to the corresponding body parts and in the same direction, whilst the shape and structure of the artificial body in view must match the shape and structure of a human body, as the illusion cannot be elicited by a block of wood (Petkova & Ehrsson, 2008; Ehrsson, 2020). The size (van der Hoort et al. 2011; Preston, 2014) and gender (Petkova et al, 2008) of the humanoid body seems to be less important, whilst the illusion also works well using the bodies of human strangers (Guterstam et al. 2015; Preston et al. 2018, 2016) and computer-generated bodies in virtual reality (Slater et al., 2010, Banakou et al., 2013).

To date, however, very few studies have explicitly examined how the full-body ownership percept is established during the full-body ownership illusion and how such a whole-body gestalt relates to the sense of ownership of specific body parts. Overall, previous research has demonstrated comparable magnitudes of subjective ownership of the artificial body irrespective of which singular body segment receives the illusion-inducing, synchronous visuotactile stimulation (Petkova & Ehrsson 2008; Petkova et al. 2011). In Gentile et al. (2015), visuotactile stimulation was applied either to the right hand, the abdomen or the right leg, and ownership of each of these three body parts was assessed. The authors observed increased ownership, not only for the specific body part that received the synchronous visuo-tactile stimulation, but also, for the other (two) non-stimulated body parts, suggesting the illusion of ownership had indeed spread to encompass the whole body; although, explicit sensations of owning the entire body were not collected in this study. In addition to questionnaire data, Petkova and Ehrsson (2008) applied visuotactile stimulation to either the right hand or the abdomen, whilst threat-evoked skin conductance responses (SCRs) (μS) to a knife, always aimed at the mannequin’s abdomen, provided an objective quantification of illusory ownership (Armel & Ramachandran, 2003). Critically, the magnitude of participants’ SCRs (μS) were not affected by whether the stimulated body part was also the one that was subsequently presented with the knife (the abdomen), or not (the hand) (Petkova & Ehrsson, 2008). Together, these behavioural insights gave weight to the hypothesis that the feeling of ownership during the perceptual illusion is not simply restricted to the local bodily site receiving synchronous multisensory stimulation, but instead becomes generalised into a global percept of ownership that is contiguous with the entire body plan. However, the precise mechanisms underlying this “spread of ownership” from the synchronously stimulated bodily site to the seamless percept for the whole body remain to be fully understood. Specifically, more research is needed to better understand the relationship between body-part and full-body ownership: is the whole simply the sum of the parts, or is the whole-body ownership experience a more complex, holistic percept that cannot be deduced entirely from its parts?

In continuation from previous studies, which only ever stimulated a singular body part at any given time, the present study set out primarily to examine the effects of stimulating multiple body parts simultaneously, as a means of potentially manipulating the illusory feeling of full-body ownership into gradations of intensity in relation to the number of body parts stimulated (1,2 or 3). For example, during a related paradigm, the invisible full-body ownership illusion (Guterstam, Abdulkarim & Ehrsson, 2015), stimulating all the invisible contours of the body plan, albeit sequentially, was beneficial in constructing the illusory percept of ownership for an entire invisible body. Likewise, perhaps when the fake body is in full view, as is the case of experiments with a mannequin’s or a stranger’s body, the volume of multisensory information congruent with the illusory percept of ownership might influence feelings of ownership for the entire artificial body. Moreover, by supplying multiple stimulations simultaneously, these signals may also be integrated within the same temporal binding window (Wallace & Stevenson, 2014; Holmes & Spence, 2004; Constantini et al. 2016), which might potentiate the resulting illusory full-body ownership percept. Therefore, for instance, it could be that converging multisensory stimulation across multiple segments of the body facilitates the illusion by increasing the amount of available perceptual evidence in support of the whole body being one’s own (Samad, Chung & Shams, 2015; Maselli & Slater, 2013; Kilteni, Maselli, Kording & Slater, 2015; Chancel & Ehrsson, 2020). However, it is also possible that a maximal illusion is elicited by the congruent visuotactile stimulation of one body segment, as, indeed, earlier studies have described a successful full-body ownership illusion by stimulating single body parts (van der Hoort, Guterstam & Ehrsson 2011; Schmalzl & Ehrsson 2011; Guterstam et al., 2015; Preston et al., 2015). No less interesting, the lack of an effect by stimulating multiple body segments simultaneously may be taken as evidence that perceived full-body ownership is not constructed simply by summation of that across constituent body parts.

In light of these unanswered questions, we conducted an experiment that, first, aimed to extend previous findings by determining whether full-body ownership can be potentiated by increasing stimulation across multiple body segments simultaneously, as compared to the stimulation of fewer or indeed a single body part. We applied the full-body ownership illusion and a within-subjects 2 x 3 design to examine the effects of synchronous versus asynchronous visuotactile stimulation involving one, two or three body parts simultaneously. For continuity with previous studies (Petkova and Ehrsson 2008; Petkova et al. 2011; Gentile et al, 2015), stimulated body parts involved (1) the trunk, (2) the trunk and the right arm, or (3) the trunk, the right arm and the right leg. The main focus of the study was to quantify the subjective experiences of body part and full-body ownership using questionnaires and to this end we developed new statements that specifically addressed ownership for five specific body parts individually (Q3 – Q7: the right arm, the left arm, the trunk, the right leg and the left leg), in addition to an explicit ownership experience for the entire body (Q8), illusory referral of touch phenomena (Q1, Q2) and the control items (Q9, Q10). With these new questionnaire items (Q3 – Q7), we aimed to investigate (1) whether increasing the numbers of stimulated body parts leads to increased full-body ownership and (2) the spread of ownership from stimulated part(s) to non-stimulated part(s) to whole. The subjective questionnaire was complemented by both threat-evoked SCR (μS) and illusion onset time (seconds) in the same participants; measures to probe the physiological and temporal dimensions of the full-body ownership illusion, respectively. Finally, inspired by recent discussions about the relationship between body ownership and interoception (Tsakiris, Tajadura-Jiménez, & Costantini, 2011; Crucianelli, Metcalf, Fotopoulou & Jenkinson, 2013; Crucianelli, Krahé, Jenkinson & Fotopoulou, 2018; Park & Blanke, 2019), we further decided to take the opportunity to explore possible links between the magnitude of the full-body ownership illusion (Q8) and individual differences in interoceptive sensitivity (Garfinkel, Seth, Barrett, Suzuki, & Critchley, 2015; Craig, 2000), as probed by the Body Awareness Questionnaire (BAQ) (Shields, Mallory & Simon, 1989).

## Methods

### Participants

48 healthy adults, a sample size that was determined before the data collection started based on previous studies (Preston et al., 2018 and Kalckert et al., 2014; both N = 40) and for the purposes of counterbalancing (see further below), were recruited to participate in the experiment via online advertisements, posters and personal communication; 28 males, 20 females, mean age 26.9 ± 6.2 years, age range: 19 - 43 years, 47 right-handed, 1 left-handed (self-reported). All had correct or corrected-to-normal vision and were instructed to wear comfortable clothing that would not interfere with the delivery of the tactile stimuli during illusion induction, i.e. no buttoned shirts or high-waisted jeans, both of which impeded the delivery of the stimuli during piloting. All recruits were naïve to the full-body ownership illusion, confirming that they had not participated in a similar study before. Participants provided written informed consent and the provided information did not explicate the purposes of this specific experiment or the details of the various experimental manipulations. The study was approved by the Swedish Ethical Review Authority (https://etikprovningsmyndigheten.se/) and conforms to the Declaration of Helsinki. After the completion of the experiment, one and a half hours in total, participants were compensated with one cinema ticket.

### Experimental set-up and illusion paradigm

Visual stimulation comprised six pre-recorded movies of a trained experimenter using custom-built, plastic hand-held probes to apply tactile stimulation to the body of a life-sized, male mannequin, which was presented from the natural (first-person) visual perspective, in an anatomically plausible and reproducible position, i.e. comfortable and supine (Fig. 1 a - c). Visual stimulation was recorded using two GoPro cameras (GoPro HERO4 Silver, GoPro Inc., San Mateo, CA, USA), mounted above a mannequin’s body so as to provide two monocular recordings from the first-person perspective, and edited using Final Cut Pro X Version 10.4.5. This software combined the two recordings so as to generate a three-dimensional, stereoscopic image of the body from the first-person perspective when presented through the head mounted display (HMD) system, for which, we used Oculus Rift DK 2 (California, USA). These steps helped to ensure that the fake body spatially substitutes one’s own as much as possible when presented in the HMDs. Half of the movies were two minutes in duration, the other half were two minutes and ten seconds. The two-minute movies were designed for the initial questionnaire session of the experiment, whilst the latter were elongated slightly for the subsequent session, including either the presentation of a knife for the SCR recording or the equivalent time as a still image, depending on the experimental condition. Another distinction between shorter and longer movies was the visual inclusion of additional equipment in the latter to obtain threat-evoked SCR (μS) and illusion onset time (seconds) data (Biopac Systems Inc., MP150; Goleta, California, USA). Specifically, two recording electrodes were seen to be attached to the middle and ring finger of the mannequin’s right hand, as they were for the participants, whilst its left hand was placed inside a covered black box containing the keypad required to indicate illusion onset time (seconds) (Fig. 1 c). During these experimental conditions, participants also placed their real left hand in the box, which was covered with black material to mask any visuo-motor incongruency induced by the real hand’s movements during button presses, which might otherwise diminish the full-body ownership illusion (Petkova & Ehrsson, 2008; Kalckert & Ehrsson, 2012; Kokkinara & Slater, 2014).

**Fig. 1.**
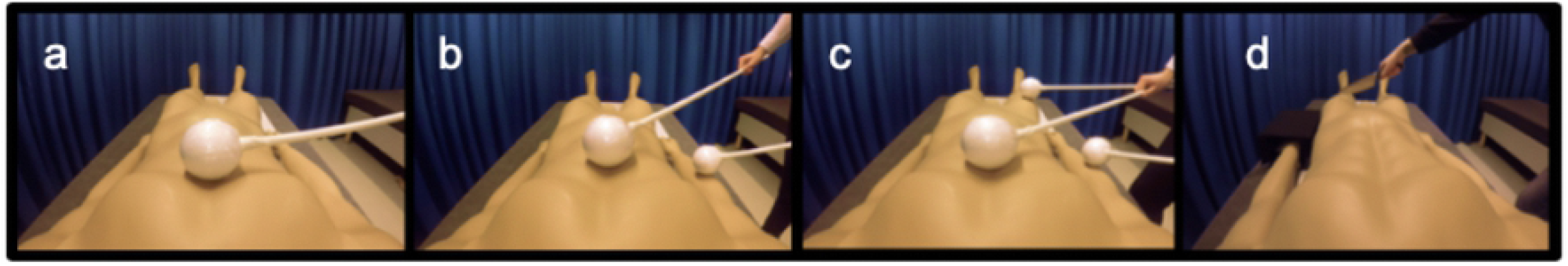
a, b, c & d. Visual stimulation in the experiment. Display of an artificial mannequin’s body from the first-person perspective and the experimenter applying tactile stimulation to one (a), two (b) or three (c) body parts simultaneously. Visual stimulation was identical for both the synchronous and asynchronous conditions, the difference being the timing of the applied strokes to the participant’s real body. The participant lay on a bed with their head tilted forward and observed these videos through a set of head mounted displays (HMDs). Panel d displays the final scene comprising the knife in the SCR and illusion onset experiment (which included only the 1S, 3S and 3A conditions; a still image was instead presented at the end of 1A, 2A and 2S). Note. Presented images appear askew as they are monocular for illustrative purposes; 3D binocular view (not askew) is only achieved within the HMDs.

Each of the spherical tactile stimuli consisted of a white, polystyrene ball with a diameter of eight centimetres, attached to a stick of one meter for the experimenter to hold. The very same probes were then used to stimulate participants’ real bodies during the experiment. Each movie, representing one of six experimental conditions, contained sixteen independent visuotactile stimulations separated by a still image of the body and surrounding scene, representing an inter-stimulus interval that ranged from four to nine seconds in duration (6.5 s on average). This frequency of visuotactile stimulation was some seconds slower than earlier studies (Petkova et al. 2008, 2011; Guterstam et al, 2015) because the longer periods of non-stimulation time were beneficial for the experimenter to accurately prepare, position and align the multiple stimuli for as close to perfect execution as possible. Tactile stimulation covered a trajectory of fifteen centimetres on the corresponding body part(s) and was always one second in duration; the onset of the first visuotactile stimulation occurring at precisely twelve seconds. For asynchronous stimulations, a two-second stimulus onset asynchrony (SOA) was introduced between the tactile stimulation(s) and the visual stimulation(s) (see further below). All the movies were identical in terms of timing; the only factors to vary were (1) the synchronicity of the visuotactile stimulation and (2) the number of stimulations occurring simultaneously. On the basis of previous studies (Gentile et al 2015), we decided that stimulated body parts would comprise the trunk (one body part), the trunk, plus the right arm (two body parts) or the trunk, plus the right arm and the right leg (three body parts) (Fig. 1 a - c). Therefore, in a within-subjects 2 (synchronicity) x 3 (number of parts stimulated) design, the six experimental conditions were: one body segment with synchronous visuo-tactile stimulation (1S), one body segment with asynchronous visuo-tactile stimulation (1A), two body segments with synchronous visuo-tactile stimulation (2S), two body segments with asynchronous visuo-tactile stimulation (2A), three body segments with synchronous visuo-tactile stimulation (3S) and three body segments with asynchronous visuo-tactile stimulation (3A).

### Procedures

Prior to commencing the illusion, participants were instructed to lie on a bed with their head tilted approximately 30 degrees forward supported by pillows and adopted a posture in which they could comfortably view their entire body. Participants then spent a few minutes adjusting the HMDs, showing only a still image of the mannequin’s body, for optimal clarity and were instructed to match their body posture to that of the mannequin as accurately and comfortably as possible, before any stimulation began. Maintaining both a comfortable and similar bodily posture facilitates body ownership illusions via visuo-proprioceptive integration, which has alone even been shown sufficient for some individuals to experience a full-body ownership illusion (Bergström et al., 2016; Carey, Crucianelli, Preston & Fotopoulou, 2019). Finally, participants were instructed to observe and attend to the whole body rather than fixating on any particular part, which may else encourage biases as a result of overt attention. Participants wore a pair of earplugs to eliminate sounds that could potentially influence the illusion experience, i.e. the sounds of the tactile stimulus touching the participants’ real bodies (Radziun & Ehrsson, 2018). After participants put on and adjusted the HMDs, inserted the earplugs and prepared a comfortable posture that matched that of the mannequin as well as possible, the experimenter initiated the movie and began applying stimulation upon carefully designed audio cues. All instructions aforementioned were repeated to the participant before each movie (i.e. experimental block) began.

Whilst participants observed the sequence of tactile stimulation being applied to the mannequin’s body with their real body occluded from view, the experimenter applied either temporally synchronous (illusion) or asynchronous (control) tactile stimulation to the participant’s real body to induce the illusion or control conditions, respectively. During conditions of multiple stimulation, the timing, force and duration were carefully controlled to match as closely as possible that witnessed by participants in the HMDs. This was achieved using carefully designed audio instructions using Audacity Version 2.2.1, which were supplied only to the experimenter via noise-cancelling headphones. The audio instructions contained auditory cues pertaining to the onset and duration of the tactile stimulation; pure tones to announce the stimulation one second before onset; white noise to indicate the duration of tactile stimulation in a vertical, downwards trajectory. These cues were overlaid on a metronome with a tempo of 120 bpm such that two beats correspond with exactly 1 second in real-time. The metronome was maintained audible in the track even during the period of white noise signalling the delivery of tactile stimulation, allowing for very precise timing. For asynchronous conditions, the onset of this audio (with respect to the movie viewed only by the participants within the HMDs) was simply delayed by two seconds, providing our SOA of two seconds with respect to the synchronous condition. To clarify, during asynchronous stimulation, the onset for the visual stimulation always preceded the tactile by precisely two seconds, ensuring no overlap (a whole one second gap) between seen and felt touches.

All participants experienced the six-condition collective three times; once for the initial questionnaire data collection, completed retrospectively at the end of each movie, and twice for the threat-evoked SCR (μS) and illusion onset time (seconds) data collection. The questionnaire session always preceded the SCR and illusion onset time sessions; since the subjective questionnaire data was the main priority, we wanted the participants as naïve as possible when completing this. To circumvent the likely influence of order effects on each of our measures, the order of the individual questionnaire items was different upon each presentation to the participant and we carefully counterbalanced the presentation of the six experimental conditions using pseudo-randomisation of synchronous and asynchronous in alternating blocks; a counterbalancing procedure that was applied to both questionnaire and SCR/illusion onset time sessions. A total of twelve different possible orders for the counterbalancing of the experimental conditions also provided the motivation to recruit a sample size of 48, affording four repetitions of each pseudo-randomisation across the participants.

### Questionnaire (first session)

The questionnaire session was designed to assess participants’ subjective experiences using a novel 10-item questionnaire formulated specifically for the purposes of the experiment (Table 1). The questionnaire was distributed immediately after each movie, statements arranged in a different order upon each presentation, always beginning with the header: “during the experiment, there were times when...” Responses were made on a 7-point Likert scale from ‘-3’ to ‘+ 3’, describing the full range of agreeability from ‘strongly disagree’ to ‘strongly agree’, where ‘0’ represents uncertainty. The questionnaire contained items pertaining to participants’ experiences of referral of touch (Q1, Q2) and illusory whole-body ownership (Q8), plus control items to assess task-compliance and suggestibility (Q9, Q10), based upon those used previously for the subjective assessment of the full-body ownership illusion (Guterstam et al., 2015; Petkova & Ehrsson, 2008). Item Q8 was considered particularly important in this study, as it represented the explicit experience of owning the entire, artificial body. We additionally formulated five body-part-specific ownership statements, referring to the illusory ownership of the mannequin’s right arm (Q3), left arm (Q4), trunk (Q5), right leg (Q6) and left leg (Q7).

**Table 1.** Questionnaire statements for the full-body ownership illusion including novel items for parts

| Item | Statement | Purpose |
| --- | --- | --- |
| Q1 | I felt the touch(es) given to the mannequin’s body | Referral of touch |
| Q2 | It seemed as though the touch(es) I felt were caused by the probe(s) touching the mannequin’s body | Referral of touch |
| Q3 | I felt as though the mannequin’s right arm were my arm | Body-part ownership |
| Q4 | I felt as though the mannequin’s left arm were my arm | Body-part ownership |
| Q5 | I felt as though the mannequin’s trunk were my trunk | Body-part ownership |
| Q6 | I felt as though the mannequin’s right leg were my leg | Body-part ownership |
| Q7 | I felt as though the mannequin’s left leg were my leg | Body-part ownership |
| Q8 | I felt as though the mannequin’s whole body were my own body | Full-body ownership |
| Q9 | I felt as though my real body were turning into a plastic body | Control |
| Q10 | I felt naked | Control |

### Threat-evoked skin conductance response (second session)

After a small break, participants commenced two repeats of the second session, allowing us to obtain threat-evoked SCRs (μS) for each of the targeted conditions: 1S, 3S and 3A. Given that SCRs are known to diminish by habituation, reduced responses resulting from repeated exposure, we chose to only present the knife within these three experimental conditions, which comprised an identical pre-recording of the experimenter presenting a large kitchen knife to the thigh region of the mannequin’s left leg (Fig. 1d) after the two minute period of either synchronous or asynchronous visuotactile stimulation, applied to one, two or three body parts simultaneously (Fig. 1a - c). In line with good ethics practise, participants were reassured that although they may witness a knife to the mannequin in some of the movies, there was never any threat to their real body during the experiment; information we also communicated clearly before participants gave their informed consent and even during the recruitment process.

SCRs (μS) were recorded continually throughout the experiment with a Biopac MP150 (Biopac Systems Inc., Goleta, USA) and registered in the accompanying software (Acqknowledge 4.9). After applying conductive electrode gel (Biopac Systems Inc., Goleta, USA) to the bottom surface of the third phalange of the index and middle finger of the right hand, the two recording electrodes were attached to the participant (Biopac Systems Inc., Goleta, USA). The left hand was positioned inside the black box containing the keypad for illusion onset time measurements (see further below). We collected the raw tonic signal at a sample rate of 100Hz and analysed the data using the same manual extraction protocol and for the same parameter of interest, the magnitude of the skin-conductance response, as that described by Petkova and Ehrsson (2008).

### Illusion onset time (second session)

In the same session as SCR recordings, we also measured the full-body illusion onset time (seconds) for conditions 1S, 2S and 3S to examine also whether the addition of stimulated body parts catalyses illusion onset. Participants placed their left hand inside the black box (Figure 4) and their left index finger over the button within, in preparation to give a single button press at their volition to indicate “the very first instance you experience the illusory sensation, so as to feel as though the mannequin’s whole body were your own body”. Participants were reminded to press the button only once per movie and to simply abstain from pressing the button if they did not specifically perceive a full-body ownership illusion. Onset times were recorded from the onset of the first visuotactile stimulation.

### Body Awareness Questionnaire

At the very end of the experiment, all participants completed the 18-item Body Awareness Questionnaire (BAQ) by Shields, Mallory and Simon (1989). This is a validated self-report scale for the measurement of individual differences in attentiveness to non-emotive, everyday bodily processes (Mehling et al., 2009), where its subscales address individuals’ attentiveness to: 1) bodily responses or changes, 2) predicting bodily reactions, 3) the sleep-wake cycle and 4) the onset of illness. The logic behind the addition of this measure was to explore the potential relationship between individual differences in ‘interoceptive sensibility’ (Garfinkel et al., 2015) and the magnitude of the full-body ownership illusion experienced by participants (Q8), reflecting an explorative attempt to account for some of the inter-individual variation that might characterise some of the range in susceptibility to the full-body ownership illusion. As the trunk of the mannequin’s body may relate to the greatest discrepancy in terms of accessible interoceptive signals, namely, its lack of breathing, it is possible that individuals most tuned to their own internal bodily processes will show reduced illusory ownership for this body part. For example, Monti et al. (2020) recently showed that illusory ownership for a virtual body could be augmented solely by using a breathing rhythm that was synchronised with participants’ actual breathing. Therefore, we also examined the correlation between participants’ self-reported BAQ scores and ownership ratings for the mannequin’s trunk (Q5).

### Statistical analysis

The majority of statistical analyses were conducted in SPSS Version 26 (IBM); however, for accessibility and availability reasons, a Bayesian paired t-test (default prior) was conducted in JASP, Version 0.9.2. All of the datasets were found to be non-normally distributed (Shapiro-Wilk test of normality) and, therefore, the appropriate non-parametric tests were utilised to investigate the effects of synchronicity and the number of body segments stimulated simultaneously (2 x 3). In the cases of multiple comparisons, we controlled for the inflated risk of incurring a Type 1 error using the Benjamini-Hochberg False Discovery Rate (BH-FDR) (McDonald, 2014), a common alternative to Bonferroni, which can be too conservative. The only situations in which we did not control for multiple comparisons was in the case of the SCR data, since we only had two planned comparisons for this data; in contrast, the questionnaire data had quite a large total number of tests (five body parts, six conditions). In general, we applied Friedman’s test to investigate whether any significant differences were present between the six experimental conditions, followed by Wilcoxon’s signed ranks tests for the scrutiny of where those significant differences lie (planned comparisons, see further below). This routine was computed individually for each of the questionnaire items, but also between experimental and control items in synchronous conditions, as well as between the control items themselves for matters of completeness (see further below).

For statistical consistency, all planned comparisons and post hoc tests involving the questionnaire data were computed according to a 2-tailed hypothesis and, where appropriate (where more than two comparisons were made), assessed using BH-FDR (FDR = 0.05) (McDonald, 2014). All p values are reported with the corrected p value (p_FDR_). We used 2-tailed tests (with the exception of the SCR analysis, see below) because, to the best of our knowledge, this study is the first that examines the effect of increasing the number of stimulated body parts in a full-body illusion paradigm, and although we had directed hypotheses in many cases, we were in interested in examining possible changes in both directions.

The questionnaire data was analysed separately for each individual illusion related (Q1-Q8) and control item (Q9, Q10) using a Friedman’s repeated-measures ANOVA followed by six pair-wise comparisons using Wilcoxon’s signed ranks tests. Three contrasts assessed the main effect of stimulation, synchronous versus asynchronous (contrasts: 1S – 1A; 2S – 2A; 3S – 3A), and three assessed the main effect of increasing the number of body parts receiving synchronous visuotactile stimulation (2S - 1S; 3S - 2S, 3S - 1S). These analyses allowed us to investigate whether illusory full-body ownership increases significantly with the addition of stimulations, but also, whether ownership for body parts, both stimulated and non-stimulated, increased significantly following the application of synchronous stimulation (Supporting Information - Table S2).

For the critical full-body ownership item, Q8, we further calculated an interaction term by comparing the difference in full-body ownership ratings between synchronous and asynchronous stimulation for the most extreme conditions, the stimulation of three body parts versus one, using the contrast [(3S - 3A) - (1S - 1A)] in a Wilcoxon’s signed ranks test. The resulting interaction term is particularly important for the focus of the current experiment, examining specifically whether the combination of synchronous stimulation and its delivery to multiple body parts simultaneously significantly facilitates the illusory percept of full-body ownership; more meaningful than a main effect of the number of stimulated body segments, per se. Post hoc, we additionally decided to analyse the illusory full-body ownership ratings between 3S - 1S in a Bayesian paired t-test (default prior). However, without the use of an informed prior (none available based on previous literature), the interpretation of Bayes Factors is debatable (Tiendo, 2019). Therefore, the result is to be interpreted with caution (Quintana & Williams, 2018).

After observing asymmetry in illusory ownership ratings for the left versus the right half of the mannequin’s body (‘hemibody’), as well as upper versus lower bodily sites, it was of interest to analyse post hoc whether the differences across the vertical and horizontal planes of the body plan were statistically significant across each synchronous condition. Using Wilcoxon’s signed ranks tests, we assessed the difference in body part ownership ratings between the stimulated right and the corresponding non-stimulated left body part.

Next, we analysed the relationships between ownership ratings for both stimulated and non-stimulated parts and full-body ownership using Spearman’s rank correlations, which gave us an indication of whether they describe related phenomena (irrespective of causality) for each of the three synchronous conditions, as well as whether the correlation co-efficient (i.e. the strength of this relationship) changes with respect to the number of stimulated body parts (1, 2 or 3). After these correlations were found to be similarly strong and significant, for both the raw ratings and when the difference between sync - async ratings were used, an ordinal regression was computed for full-body ownership ratings with body part ownership ratings as the predictor variable, which was conducted separately for all parts, stimulated parts only and non-stimulated parts only (post hoc). After also checking that the number of synchronous stimulations (1, 2 or 3) was indeed an irrelevant manipulation by computing an ordinal regression for full-body ownership ratings by the number of synchronous stimulations, we included all synchronous conditions’ data rather than running the analysis separately for 1S, 2S and 3S.

Finally, for the control analyses, ratings to the control items (Q9, Q10) were compared against ratings for the experimental questionnaire items (Q1-Q8) using Wilcoxon’s signed ranks. Significant differences between these variables is a complementary validation that the subjects’ responses to the latter reflect their experience of the illusion as opposed to mere confabulation or task compliance, thus reducing the likelihood of demand characteristics.

Threat-evoked SCR data (μS) was also analysed using Wilcoxon’s signed ranks tests. Our two planned comparisons were designed to compare only the magnitude of the threat-evoked SCR (μS) between synchronous and asynchronous visuotactile stimulation, 3S - 3A (one-tailed), to assess the basic effect of the illusion, and 1S - 3S (two-tailed), to examine the effect of increasing the number of synchronously stimulated body parts (from one to three simultaneously). Prior to performing the tests, an average response for each participant was calculated separately for each condition. Data from two participants was excluded due to technical issues during signal acquisition. For the remaining 43 participants, SCRs (μS) were identified as the peak-to-peak magnitude of the very first waveform (Braithwaite, Watson, Jones & Rowe, 2013) to follow the onset of the knife threat, which, to reiterate, was always presented to the mannequin’s left leg (maximum 7 seconds post onset of the threatening stimulus). Trigger codes ensured these epochs were identifiable during the offline analysis, in which, we collated and averaged both trials for each experimental condition prior to performing statistical analyses. At this stage, three data sets were removed since they contained abnormally large values; an average SCR magnitude > 4.0 μS (Braithwaite et al., 2013) and hence, N = 43 for the final analysis. We did not control for multiple comparisons in these specific analyses since the number of planned comparisons were small (two) and we had strong a priori hypothesis to expect the weakest SCR in the asynchronous condition (one-tailed: 3S versus 3A) (Petkova & Ehrsson, 2008; Preston et al., 2015; Guterstam et al., 2015). Post hoc, we extended this to the unplanned contrast, 1S - 3A (see Results for details).

Illusion onset time (seconds) for 1S, 2S and 3S was analysed in a one-way repeated-measures Friedman’s test followed by Wilcoxon’s signed ranks. Here, we were interested to explore whether the addition of stimulated body segments decreased the rate at which an explicit illusory whole-body percept emerges. However, due to the response rate of 69%, only 33 participants’ data were included in the analysis for illusion onset time (seconds). In final series of post hoc tests, we correlated the magnitude of the full-body ownership illusion (Q8) with the onset time (seconds) for 1S, 2S and 3S to examine whether there was any significant relationship between the strength of the rated illusion and its reported onset.

Finally, BAQ (Shields et al., 1989) scores were computed for each individual participant as the total of the ratings to each of the 18 items. Spearman’s rank correlations were then used in order to explore whether individual differences in the magnitude of this self-report measure of interoceptive sensibility could be related generally to the magnitude of subjective full-body ownership illusion ratings (Q8), illusory ownership of the mannequin’s trunk (Q5), threat-evoked SCRs (μS), or the rate of illusion onset (seconds). We attempted these analyses on both the synchronous data and on the difference between synchronous and asynchronous data where possible (e.g. illusion onset times were not collected in the asynchronous conditions). Many more exploratory correlations were attempted, and these are reported in the Supporting Information Fig. S2, including those using averages of the BAQ subscales (Shields, Mallory & Simon, 1989) in place of the total BAQ score. These results was also largely negative and consistent with the negative findings from the analysis with the total BAQ score (see Supporting Information, Fig. S2).

## Results

### Subjective questionnaire data: overview

For descriptive purposes and to maintain consistency with earlier studies (Petkova & Ehrsson, 2008), mean ratings for the referral of touch (Q1, Q2), illusory ownership of individual body parts (Q3 - Q7), illusory ownership of the entire artificial body (Q8) and the control items (Q9, Q10), after synchronous (illusion) and asynchronous (control) visuotactile stimulation applied to one, two or three body parts simultaneously, are presented below in Fig. 2.

**Fig. 2.**
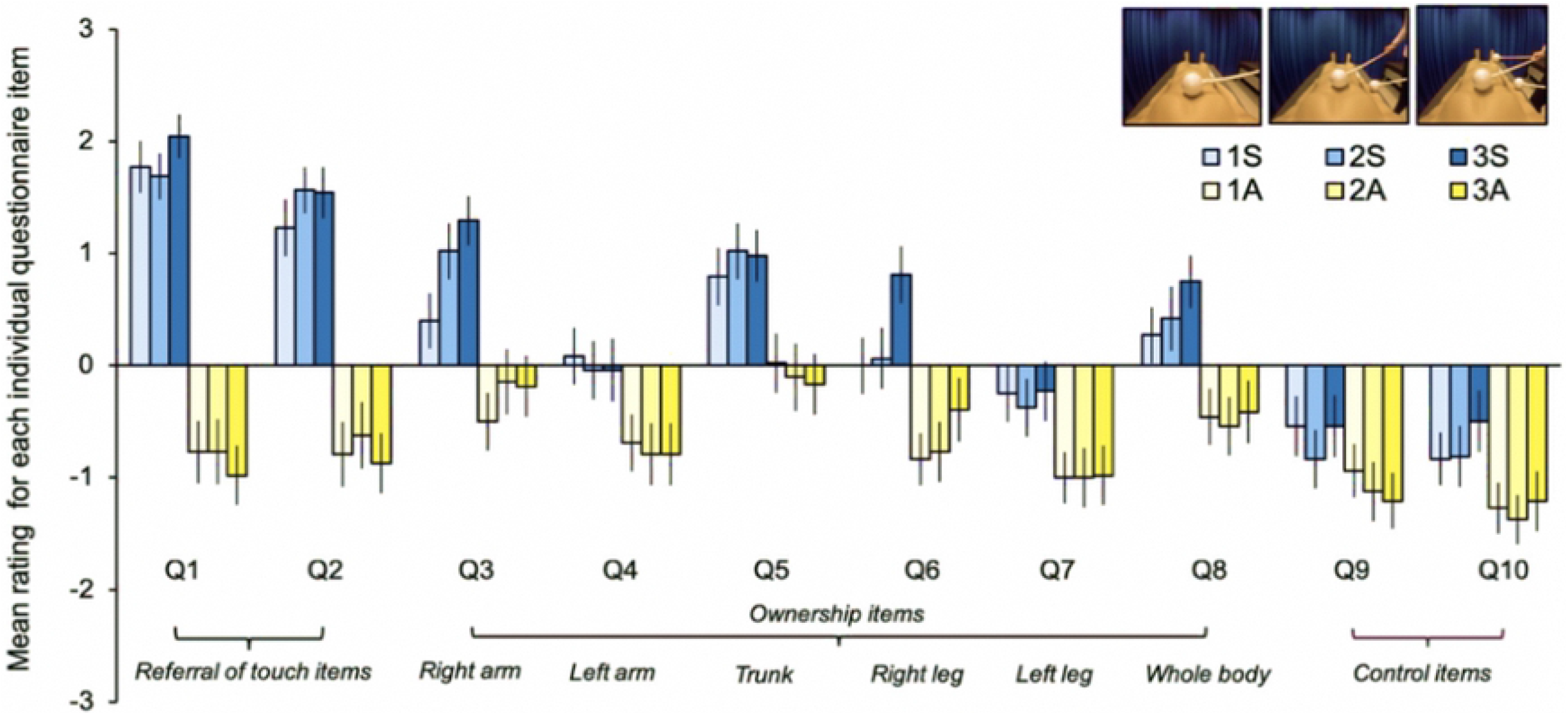
Mean ratings for each individual questionnaire item (Q1-Q10) across all six conditions. N = 48. Mean response to each questionnaire item, described by annotations within the figure, for conditions involving synchronous (blue) or asynchronous (yellow) visuo-tactile stimulation applied to one (lightest), two (intermediary) or three (darkest) body segments simultaneously. Error bars represent standard error of the mean (SEM). Presented for illustrative purposes and for comparisons with earlier studies (Petkova & Ehrsson, 2008).

On average, the majority of experimental questionnaire items (Q1-Q8), assessing referral of touch and ownership for individual body parts and the body whole, were affirmed (response > 0) by participants following synchronous stimulation, whilst they were rejected (response < 0) following asynchronous (all p values pertaining to comparisons between the synchronous and its asynchronous counterpart were significant to at least p <.01 after applying BH-FDR). All the data for the planned comparisons is presented in Supporting Information - Table S1 with the p_FDR_ and a measure of effect size, r = Z/✓N (Rosenthal, 1994). The table further contains the data for the analyses comparing control and experimental items, as well as between the two control items themselves (Supporting Information - Table S1).

### Full-body ownership does not increase significantly with the number of stimulated body parts

Results for Q8 (Table S2; Q8), the critical item in the questionnaire, referring specifically to the extent to which participants agreed with the statement “I felt as though the mannequin’s whole body were my own body”, are displayed in Fig. 3.

**Fig. 3.**
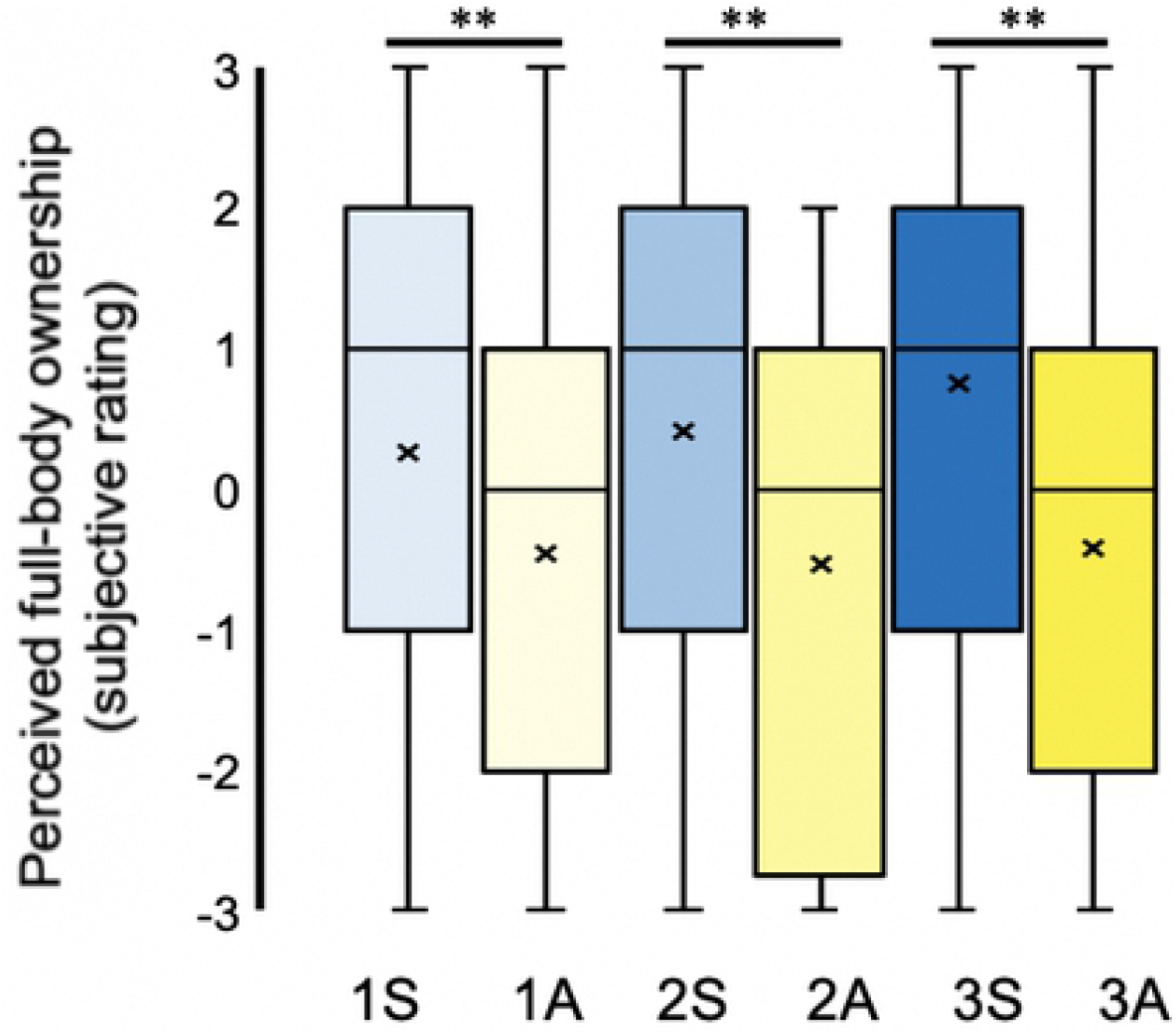
Illusory ownership for the mannequin’s whole body. N = 48. Perceived ownership for the entire artificial body across both synchronous (blue) and asynchronous (yellow) visuotactile stimulation to one (lightest), two (intermediary) and three (darkest) body segments simultaneously. The mean and median values are represented by the · and the straight line within the boxplot, respectively. Note. *** indicates significant to p <.001, ** to p <.01 and * to p <.05 after Benjamini-Hochberg FDR correction.

First, we conducted a test for the presence of a significant difference between stimulation synchronicity and the number of body parts stimulated simultaneously [(3S - 3A) - (1S - 1A)]. This interaction term is especially important since it describes the combined benefit of applying synchronous stimulation and to multiple body parts. Indeed, this interaction effect was found to be non-significant (Z = 1.651, p =.099). Second, we conducted a Bayesian paired test (3S - 1S) to obtain a Bayes Factor (BF_10_) of 1.872 (max BF_10_: 2.678 at r =.23). This indicates that evidence in favour of the alternate hypothesis is anecdotal, being only 1.872 times greater than that of the null; generally regarded as being below the arbitrary recommended cut-off for rejecting the null hypothesis (Quintana & Williams, 2018). Third, for completeness, an ordinal regression failed to support any significant relationship between the number of synchronously stimulated body parts and full-body ownership ratings, χ2 (2, *N* = 48) = 2.052, p =.358, pseudo R^2^ (McFadden) =.004. Therefore, illusory full-body ownership may not be expressed as a linear function of the number of synchronously stimulated parts. Altogether, we regard these findings as multiple lines of evidence for a lack of significant facilitatory effect of increasing the number of body segments receiving synchronous multisensory stimulation to the subjective magnitude of the full-body ownership illusion.

### Stronger body part ownership following synchronous visuotactile stimulation

As expected, synchronous visuotactile stimulation of the mannequin’s body resulted in significantly increased ownership ratings for all individual body parts as compared to its asynchronous stimulation (Fig. 4 – 6). Specifically, this included both the parts that received the visuotactile stimulation during particular experimental conditions (trunk, right hand and right leg), but interestingly, also the left limbs, which were never stimulated. Therefore, this may be taken as evidence for the “spread of ownership” (Petkova & Ehrsson, 2008) from stimulated to non-stimulated body parts in conditions involving the delivery of synchronous visuotactile stimulation. Multisensory integration enhances illusory ownership directly in the case of the stimulated body parts, but also indirectly, as in the case of the non-stimulated body parts. The results of a regression analysis further revealed that the ratings of non-stimulated body parts (synchronous - asynchronous) could be predicted in part by the ratings of ownership for stimulated body parts (synchronous - asynchronous); χ2(25, *N* = 48) = 98.366, p <.001, pseudo R^2^ (McFadden) =.159. However, only 15.9% of the variance in non-stimulated body part ownership may be explained by variance in ownership for stimulated parts.

**Fig. 4.**
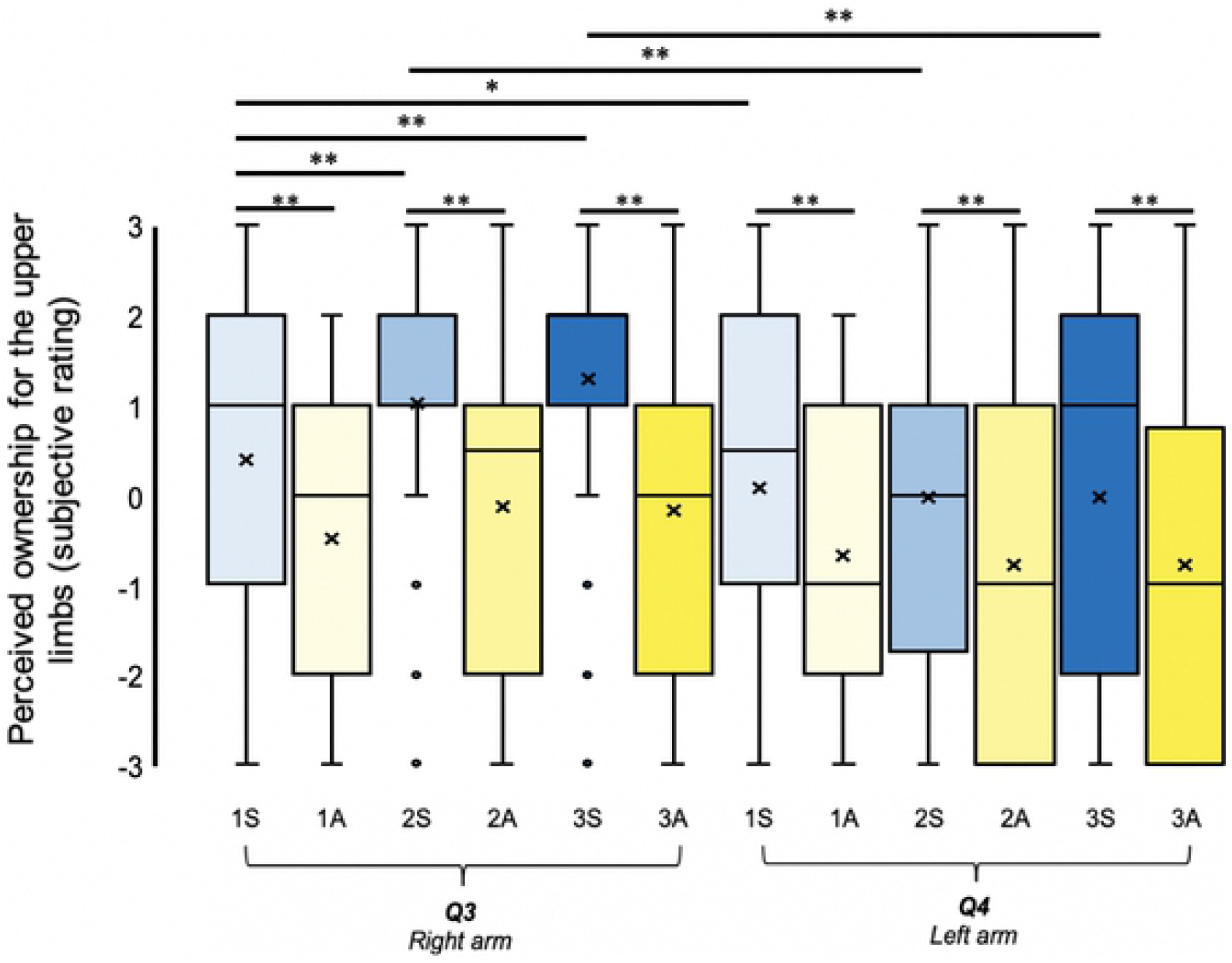
Illusory ownership for the mannequin’s right and left arms. N = 48. Perceived ownership for right arm (Q3) and left arm (Q4) across both synchronous (blue) and asynchronous (yellow) visuotactile stimulation to one (lightest), two (intermediary) and three (darkest) body segments simultaneously. The mean and median values are represented by the x and the straight line within the boxplot, respectively. Note. *** indicates significant to p <.001, ** to p <.01 and * to p <.05 after Benjamini-Hochberg FDR correction.

**Fig. 5.**
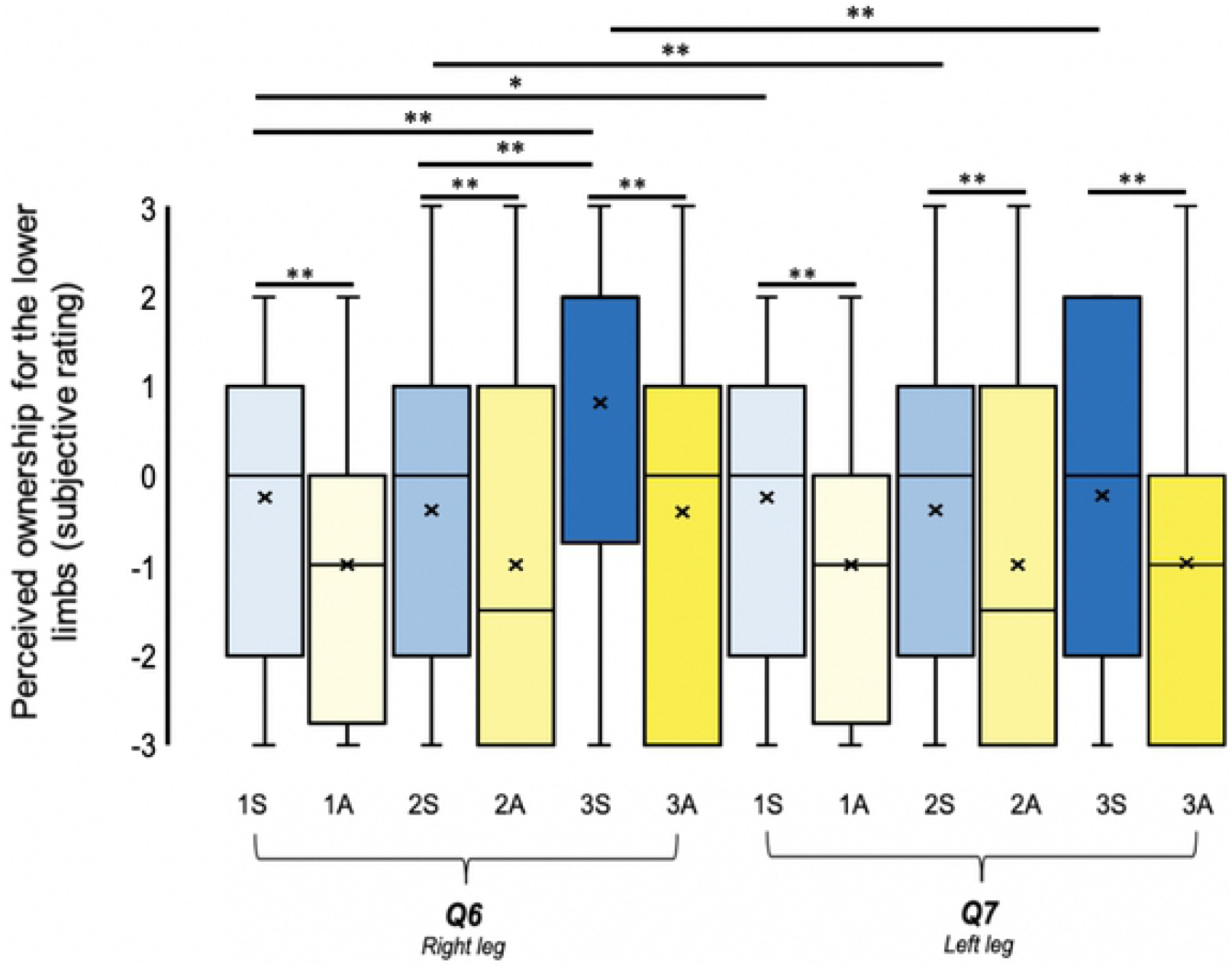
Illusory ownership for the mannequin’s right and left legs. N = 48. Perceived ownership for the right leg Q6 and left leg Q7 across both synchronous (blue) and asynchronous (yellow) visuo-tactile stimulation to one (lightest), two (intermediary) and three (darkest) body segments simultaneously. The mean and median values are represented by the x and the straight line within the boxplot, respectively. Note. *** indicates significant to p <.001, ** to p <.01 and * to p <.05 after Benjamini-Hochberg FDR correction.

**Fig. 6.**
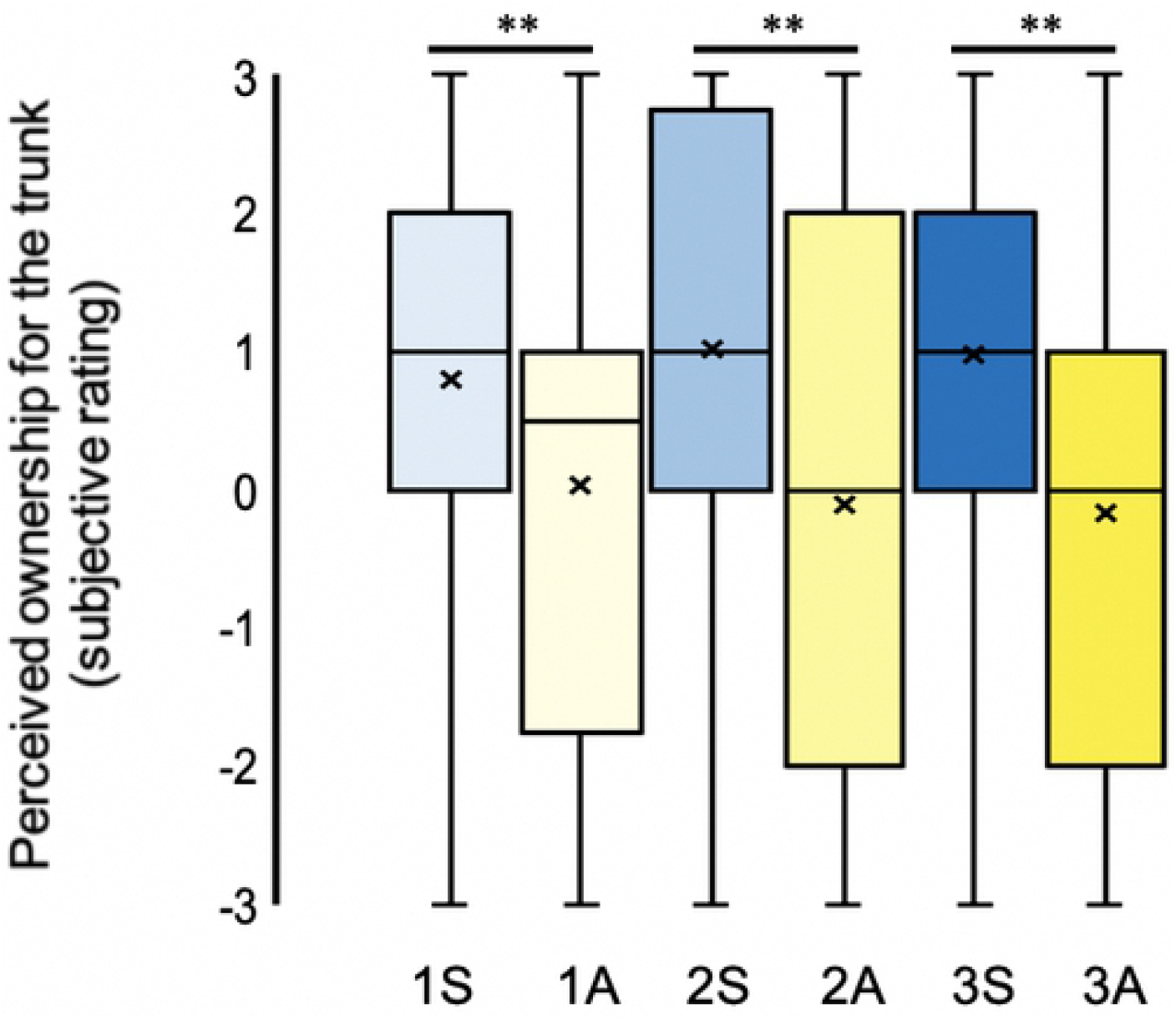
Illusory ownership for the mannequin’s trunk. N = 48. Perceived ownership for the trunk, Q5 across both synchronous (blue) and asynchronous (yellow) visuotactile stimulation to one (lightest), two (intermediary) and three (darkest) body segments simultaneously. The mean and median values are represented by the x and the straight line within the boxplot, respectively. Note. *** indicates significant to p <.001, ** to p <.01 and * to p <.05 after Benjamini-Hochberg FDR correction.

### Body parts receiving synchronous visuotactile stimulation are perceived with stronger illusory ownership than are body parts receiving no stimulation

As expected, we found rated ownership for the individual body parts to be significantly increased when they were directly stimulated synchronously, as compared to when the same body part received no visuotactile stimulation (Fig. 4 – 6), in line with Gentile et al. (2015). For example, in both 2S and 3S, conditions in which we stimulated the right arm, illusory ownership ratings for the mannequin’s right arm were significantly higher than during 1S, in which the right arm received no stimulation (Fig. 4, Table S1; Q3). Similarly, for 3S, in which we stimulated the right leg, illusory ownership ratings pertaining to the mannequin’s right leg were also rated much higher than for 2S and 1S, conditions in which the right leg received no stimulation (Fig. 5, Table S1; Q6). These findings support our hypothesis that congruent visuo-tactile stimulation boosts body ownership for stimulated parts. Consistent with this, no significant changes were observed for illusory ownership of the mannequin’s trunk; the only body segment to consistently receive synchronous stimulation (Fig. 6, Table S1; Q5).

### Subjective illusory body part ownership for the left-sided versus the right-sided limbs

For the left-sided limbs, we found similar levels of body part ownership for synchronous visuo-tactile stimulation applied to one, two or three body parts simultaneously (Table S1; Q4, Q7). However, we also had the novel opportunity to directly compare the ownership illusion for the left versus the right hemibody, which became of interest after observing consistently non-uniform illusory ownership ratings for individual body parts. In the first of these post hoc tests, regarding the upper limbs (Fig. 4), we found significant differences between the magnitude of participants’ perceived ownership for the mannequin’s left versus right arm for synchronous conditions of visuo-tactile stimulation applied to one, two or three body segments simultaneously, χ2(2) = 57.332, p <.001. Intriguingly, Wilcoxon’s signed ranks tests revealed that illusory ownership of the mannequin’s right arm was consistently perceived to a greater extent than that of the left arm, both when neither limb was receiving visuo-tactile stimulation (1S: Z = 2.354, p =.019, p_FDR_ =.019, r =.34) and, more expectedly, when the right arm received stimulation (2S: Z = 4.533, p <.001, p_FDR_ =.0015, r =.65 and 3S: Z = 4.692, p <.001, p_FDR_ =.0015, r =.68). Likewise, for the lower limbs (Fig. 5), participants rated illusory ownership to significantly different degrees across the three synchronous conditions, χ2(2) = 32.846, p <.001. Significantly enhanced ownership for the mannequin’s right leg as compared to the left was apparent, interestingly, when neither leg was stimulated (1S; Z = 1.983, p =.047, p_FDR_ =.047, r =.29 and 2S; Z = 2.946, p =.003, p_FDR_ =.0045, r =.43) and, more expectedly, when the right leg was stimulated (3S; Z = 4.692, p <.001, p_FDR_ =.003, r =.68). Therefore, left-right asymmetry was present across all loads of multisensory stimulation (1S, 2S and 3S) and for both limb types.

### Subjective illusory body part ownership for upper versus lower limbs

We also explored post hoc possible variations in illusory body-part ownership between upper and lower limbs. The results of the Wilcoxon’s signed ranks tests revealed that the upper limbs were always perceived with significantly greater illusory ownership than were the mannequin’s lower limbs; 1S: Z = 3.170, p =.002, p_FDR_ =.003, r =.46; 2S: Z = 3.475, p =.001, p_FDR_ =.003, r =.50; S3: Z = 2.735, p =.006, p_FDR_ =.006, r =.39. Next, we examined the differences in the upper versus lower limbs’ perceived ownership split by hemibody. For the right-sided arm and leg, the body parts that often received visuo-tactile stimulation, we observed a significant difference across our three synchronous conditions (right hemi-body: χ2(2) = 52.949, p <.001). Wilcoxon’s signed ranks tests further revealed that participants reported significantly greater perceived ownership for the mannequin’s right arm as compared to the mannequin’s right leg, interestingly, both when neither were stimulated (1S: Z = 2.564, p =.01, p_FDR_ =.01, r =.40), when only the right arm was stimulated (2S: Z = 4.550, p <.001, p_FDR_ =.003, r =.66) and when both upper and lower limbs were stimulated (3S: Z = 2.773, p =.009, p_FDR_ =.01, r =.40). However, a Freidman’s test revealed no significant differences in upper versus lower limb ownership ratings for the left-sided parts, which did not receive visuotactile stimulation (χ2(2) = 8.677, p =.123). Moreover, significantly increased ownership for the mannequin’s left arm relative to the left leg for 1S did not survive correction for multiple comparisons (Z = 2.560, p =.03, p_FDR_ =.09, r =.37) whilst, during 2S and 3S, conditions in which rightward bodily sites received synchronous visuotactile stimulation, no significant differences were found between illusory ownership of the left arm versus left leg (Z = 2.090, p =.06, p_FDR_ =.09, r =.30 and Z = 1.555, p =.120, p_FDR_ =.120, r =.22, respectively). Thus, there were some differences in illusion strength between arms and legs (with a stronger illusion for the arm), but this effect was only present significantly for the right (often stimulated) side of the body after correction for multiple comparisons in the current paradigm.

### Part-to-whole ownership relationships within the full-body ownership illusion

Consistently, significant, strong positive correlations were identified between rated ownership for the body part(s) receiving synchronous visuotactile stimulation (calculated as an average ratings of the relevant questions for each condition; 1S = Q5; S2 = (Q3+Q5)/2; 3S = (Q3+Q5+Q6)/3) and that of the whole artificial body (Q8); 1S: r_s_ =.68, p <.001,2S: r_s_ =.73, p <.001 and 3S: r_s_ =.85, p <.001. Therefore, the greater the illusory ownership for the stimulated body part(s), the greater the illusory ownership for the whole body. However, similarly strong positive correlations were identified between ownership ratings for the non-stimulated body part(s) (1S = (Q3+Q4+Q6+Q7)/4; 2S = (Q4+Q6+Q7)/3; 3S = (Q4+Q7)/2) and the whole body (Q8); 1S: rs =.68, p <.001; 2S: r_s_ =.70, p <.001 and 3S: r_s_ =.79, p <.001 (Fig. 7 a – c, respectively). These correlations held even when difference ratings, synchronous ratings (or the average of synchronous ratings for multiple body parts) minus asynchronous ratings (or the average of asynchronous ratings for multiple body parts), were utilised instead; 1S: (stimulated) rs =.54, p <.001, (non-stimulated) rs =.57, p <.001; 2S: (stimulated) rs =.64, p <.001, (non-stimulated) rs =.69, p <.001; S3: (stimulated) rs =.73, p <.001, (non-stimulated) rs =.53, p <.001.

**Fig. 7.**
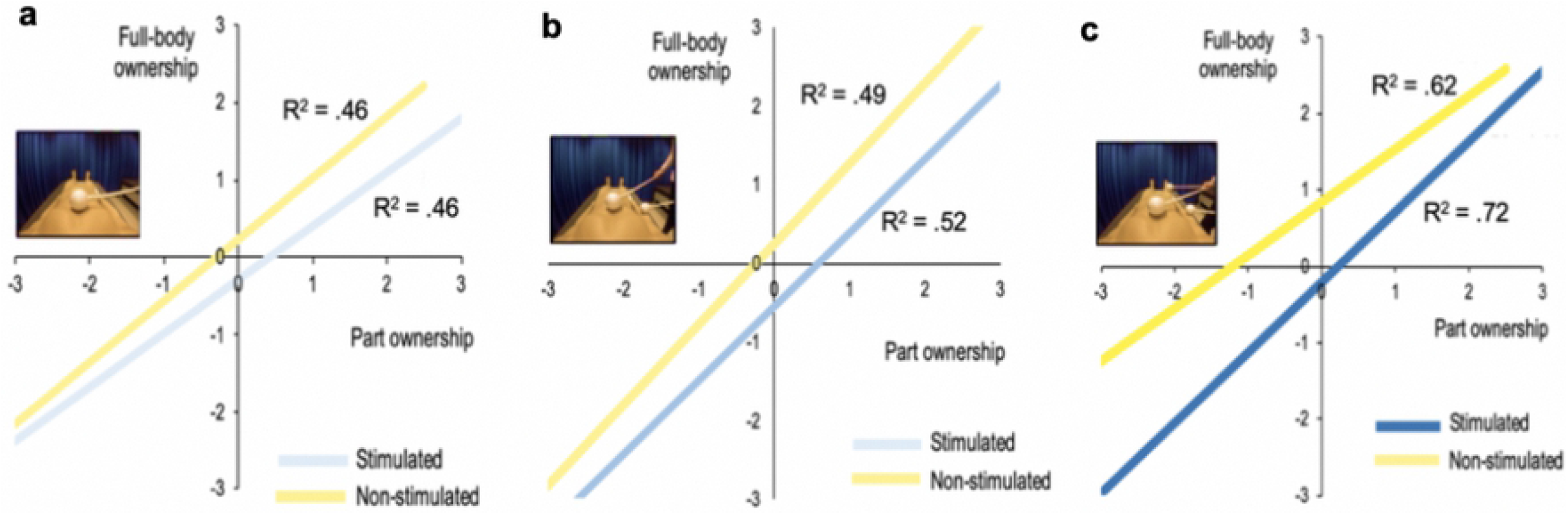
a – c. Rated illusory ownership for the whole body versus stimulated and non-stimulated parts in 1S, 2S, 3S. N = 48. Positive linear relationships between subjective ownership for the entire, artificial body and that of its parts, both synchronously stimulated part(s) (blue) (1S: trunk; 2S: trunk and right arm; 3S: trunk, right arm and right leg) and non-stimulated (neither synchronous nor asynchronous) parts (yellow) (1S: right arm, left arm, right leg and left leg; 2S: left arm, left leg and right leg; 3S: left arm and left leg. Conditions 1S, 2S and 3S are represented by Figure 7 a, b and c, respectively. In the cases of multiple body parts, an average rating was formed for their comparison with illusory full-body ownership (singular item, Q8). Data correspond to the ratings for the synchronous only (not the difference between sync-async). All correlations were found to be significant to p <.001. Concatenating condition type (1S, 2S, 3S) and subtracting the corresponding asynchronous ratings (sync-async), regression analyses for parts, stimulated parts and non-stimulated parts were also significant to at least p <.01.

In light of these results, we also computed ordinal regression analyses on the synchronous minus asynchronous ratings in order to investigate whether illusory ownership for body parts (all, stimulated and non-stimulated) predicts the magnitude of the resulting full-body ownership illusion. These regression analyses were significant in all cases, concatenating over 1S, 2S and 3S: average ratings for all parts to whole, χ2(31, *N* = 48) = 284.360, p <.001, pseudo R^2^ (McFadden) =.545 (shared variance, 54.5%); average ratings for stimulated parts to whole, χ2(15, *N* = 48) = 175.115, p <.001, pseudo R^2^ (McFadden) =.336 (shared variance, 33.6%); average ratings for non-stimulated parts to whole, χ2(26, *N* = 48) = 105.189, p <.001, pseudo R^2^ (McFadden) =.202 (shared variance, 20.2%). This suggests that the greater the feeling of ownership for stimulated parts and the greater the spread of illusory ownership to non-stimulated body parts, the greater the resultant full-body ownership percept during the illusion. This finding resonates the notion that, despite there being no beneficial effects of increasing the number of body parts receiving synchronous visuotactile stimulation simultaneously (regression using number of synchronous stimuli as a regressor on full-body ownership ratings, p =.358; see above), the magnitude of illusory ownership for the individual body segments does appear to reflect a causal relationship with the illusory percept of full-body ownership.

### Threat-evoked skin conductance response (μS)

Oddly, a statistically significant difference was found between the SCRs (μS) collected for 3S - 1S (two-tailed: Z = −2.137, p =.033, r =.33), but in the direction of responses being significantly greater for 1S than for 3S (Fig. 8). Against our expectations and against the questionnaire results described above, we also did not observe significantly stronger SCRs (μS) in the condition with synchronous visuo-tactile stimulation compared to the condition with asynchronous visuo-tactile stimulation (one-tailed: 3S - 3A: Z =.537, p =.296, r =.08). For a ‘sanity check’, we further analysed the contrast 1S - 3A post hoc, but also one-tailed, due to our strong a priori hypothesis regarding the direction of any difference and because we have used the 1S condition before (e.g. Petkova & Ehrsson, 2008). This analysis did reveal a significant difference in the expected direction (Z = 2.985, p =.002, r =.53). Thus, we conclude that the current SCR results (Fig. 8) are somewhat inconclusive and have produced only mix evidence in support of successful full-body ownership illusion (see Discussion).

**Fig. 8.**
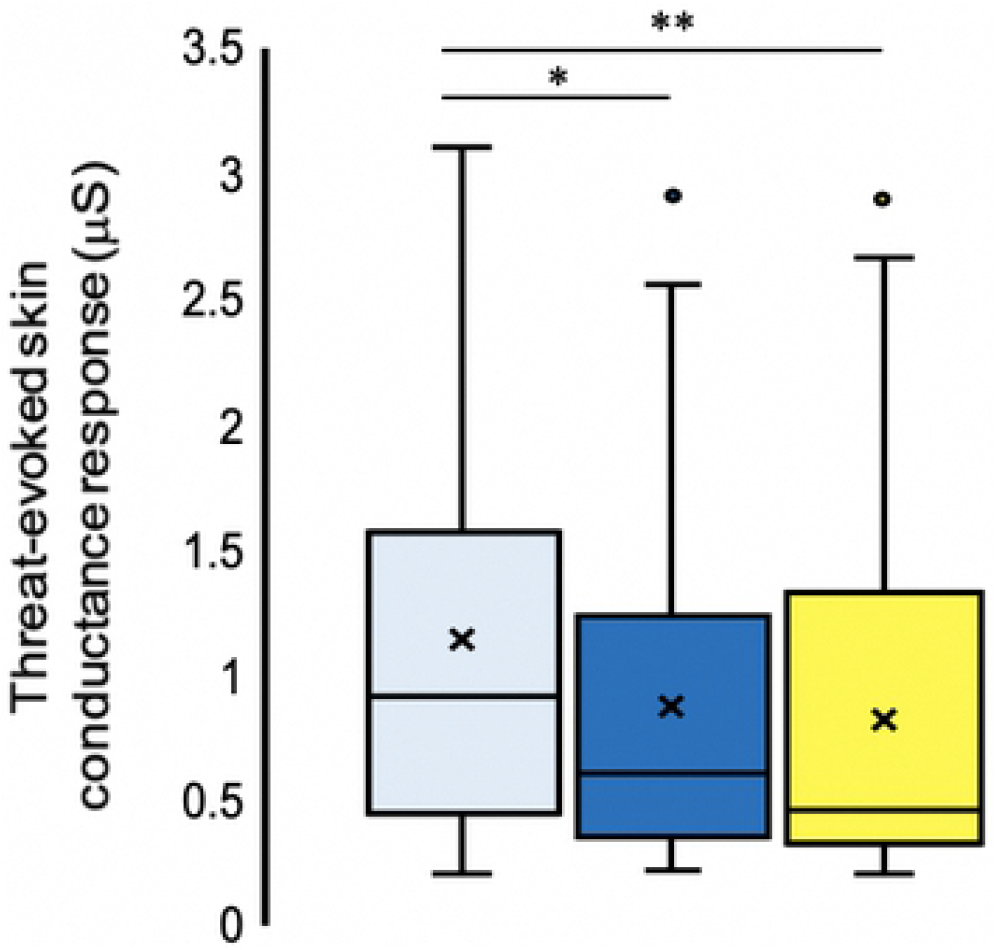
Threat-evoked skin conductance responses (μS). N = 43. Threat-evoked SCRs following conditions of synchronous visuotactile stimulation applied to three (dark blue), one body segment (light blue), and asynchronous (control) stimulation to three body segments (yellow). The mean and median values are represented by the x and the straight line of the boxplot, respectively.

### Illusion onset time (seconds)

In addition to the magnitude of the illusion, we speculated that perhaps the more body parts receiving synchronous stimulation, the faster the percept of full-body ownership should emerge. Therefore, we collected participants’ self-reported (via button-press) time to indicate “the very first instance you experience the illusory sensation, so as to feel as though the mannequin’s whole body were your own body”. We first analysed the data from the 33 participants that pressed the button at least once per experimental condition of interest (1S, 2S and 3S), and secondly the data from consistent responders, who supplied responses on both runs (N = 20). Moreover, the response rate of 69% (33/48) is roughly akin to that reported for the rubber hand illusion in Kalckert and Ehrsson (2017), around 70 - 75% of a recruited sample. Onset times for 1S, 2S and 3S were analysed using a Friedman’s test, which returned no evidence of a significant difference (N = 33: χ2(2) = 0.424, p =.809; N = 20: *χ2(2)* = 1.3, p =.522). Similarly, Wilcoxon’s signed ranks tests revealed that the time to the onset of an explicit full-body ownership illusion was unaffected by the number of body parts in receipt of synchronous stimulation (N = 33: 2S - 1S: Z = 0.688, p =.492, p_FDR_ =.611, r =.12; 3S - 2S: Z = 0.742, p =.458, p_FDR_ =.611, r =.13; 3S - 1S: Z = 0.509, p =.611, p_FDR_ =.611, r =.09 and for N = 20: 2S - 1S: Z = 1.269, p =.204, p_FDR_ =.306, r =.28; 3S - 2S: Z =.597, p =.550, p_FDR_ =.550, r =.13; 3S - 1S: Z = 1.68, p =.093, p_FDR_ =.279, r =.38). The findings (N = 33) are summarised in Fig. 9.

**Fig. 9.**
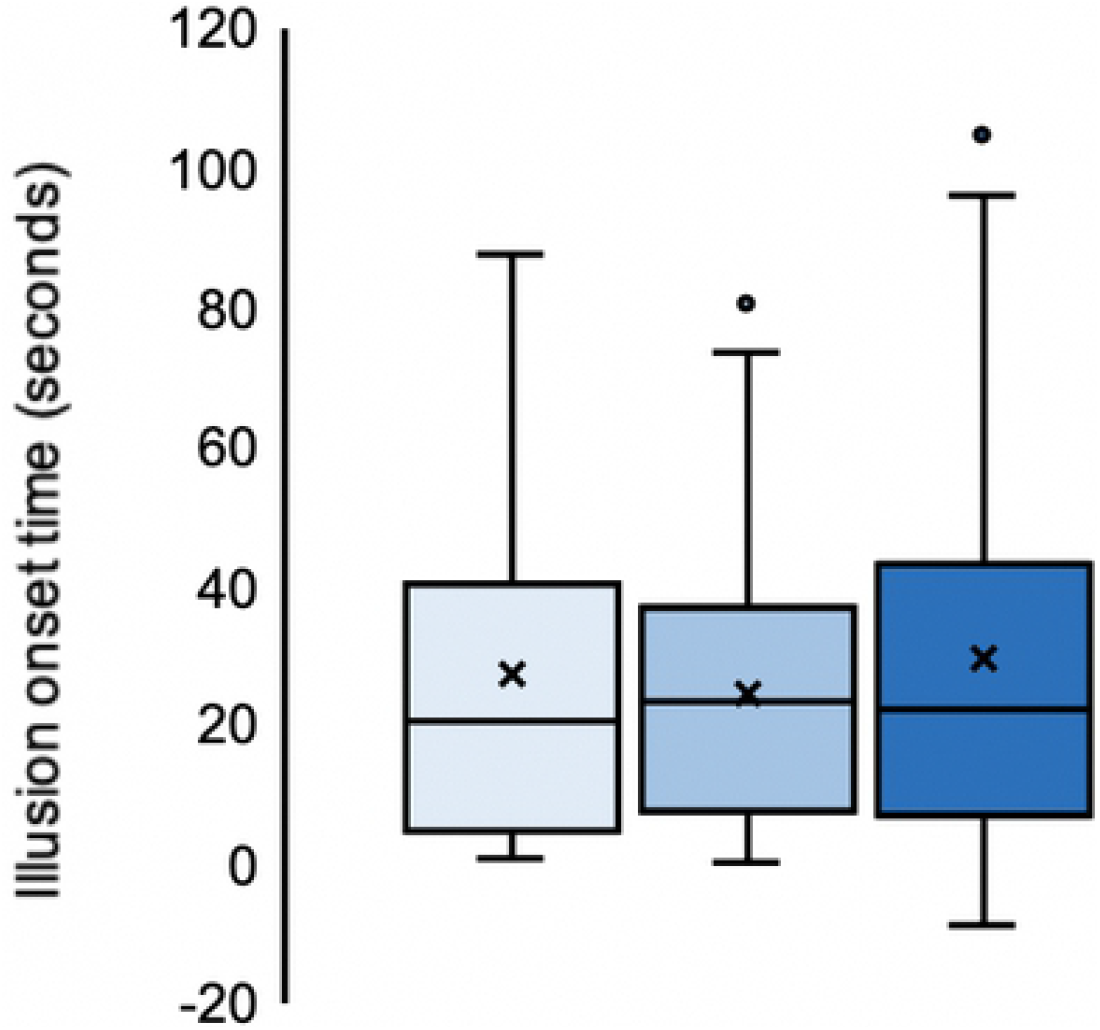
Illusion onset time (seconds). N = 33. Illusion onset times (seconds) averaged over two repeats for each two-minute stimulation of conditions involving synchronous visuotactile stimulation to one, two or three body segments simultaneously (lightest - darkest). The mean and median values are represented by the x and the straight line within the boxplot, respectively. For clarity and rounded to the nearest second: 1S: mean = 28 seconds, median = 21 seconds; 2S: mean = 25 seconds, median = 24 seconds and 3S: mean = 30 seconds, median = 22 seconds. T_0_ = the onset of the very first visuotactile stimulation, which was preceded by 12 seconds of visuo-proprioceptive stimulation as the experimenter prepared to apply the stimulations to the participants’ real body.

Illusion onset times (seconds) were also correlated with full-body ownership ratings post hoc in order to investigate the link, if any, between subjective magnitude of the illusion and the rate to onset. For clarity, this analysis was conducted only on data from the consistent responders (N = 20). Whilst Spearman’s rank correlations were non-significant for 1S (r_s_ = -.280, p =.232) and 2S (r_s_ = -.084, p =.725), for 3S, subjective illusion magnitude was found to significantly, negatively correlate with the rate of onset (3S: r_s_ = -.516, p =.02), indicating that the more participants perceived full-body ownership, the faster the onset of the illusion for 3S. However, using BH-FDR, this result might be a false positive (p_FDR_ =.06). Thus, taken together, our results are inconclusive with respect to a systematic relationship between illusion onset time and the magnitude of subjective full-body ownership illusion.

### Body Awareness Questionnaire and susceptibility to a full-body ownership illusion

Lastly, we explored how participants’ susceptibility to the full-body ownership illusion may have related to self-reported interoceptive awareness (‘interoceptive sensibility’; Garfinkel et al., 2015) of everyday bodily processes, covering awareness of responses or changes in body processes, predicting body reactions, the slee-pwake cycle and the onset of illness). To this end, we used the Body Awareness Questionnaire (Shields, Mallory & Simon, 1989). Self-rated BAQ scores, the total sum to all 18 items (mean = 4.13, SD = 0.74, SEM = 0.11), were found not to correlate significantly with rated full-body ownership, trunk ownership (and both but as the difference between synchronous and asynchronous ratings), illusion onset time nor threat-evoked SCR (see Supporting Information - Table S2). In failing to find any significant correlations, we conclude that self-reported interoceptive sensibility probably does not reflect individual variation of any relevance to the experience of a full-body ownership illusion induced by visuotactile stimulation.

## Discussion

The present experiment set out primarily to investigate whether simply increasing the number of body parts in receipt of synchronous visuotactile stimulation significantly increases perceived full-body ownership during a full-body ownership illusion (Petkova & Ehrsson, 2008). Should increasing the number of synchronously, simultaneously stimulated body parts contribute significant enhancements to the holistic whole-body ownership percept, it could be argued that the illusory feeling of full-body ownership reflects the summation of perceived ownership across constituent body parts, in turn, induced by their synchronous visuotactile stimulation(s). Using a novel questionnaire designed specifically for the purposes of analysing the explicit subjective sensation of illusory full-body ownership (Q8), but also, that of all individual body parts (Q3-Q7), we first hypothesised to replicate and extend the difference between synchronous and asynchronous visuotactile stimulation (Petkova & Ehrsson, 2008). Continuing from previous research, measures of illusory ownership for the entire artificial body, the stimulated body parts, but also, the non-stimulated body parts (including those never stimulated; the left limbs) were significantly greater following synchronous, as opposed to asynchronous, visuotactile stimulation. This was evident from the subjective questionnaire data in participants’ responses toward the experimental items that asked participants to rate the extent to which they specifically experienced the entire mannequin’s body (Q8) and each of its major constituent parts (Q3-Q7) as their own. Therefore, in addition to a full-body ownership effect, the illusion gave rise to perceptions of ownership for all of the constituent body parts. Furthermore, ownership ratings for the stimulated body parts was found to significantly predict those of non-stimulated body parts; strong evidence for the “spread” of illusory ownership between stimulated and non-stimulated parts. Regression analyses also supported ratings of ownership for both stimulated and non-stimulated body parts in predicting the ownership illusion for the entire artificial body. However, despite these findings, multiple lines of evidence, both frequentist and Bayesian, suggested that illusory full body ownership (Q8) may not be significantly enhanced by converging synchronous multisensory stimulation across multiple segments of the body simultaneously. Therefore, the magnitude of subjective full-body ownership may not necessarily, directly depend upon the volume of synchronous multisensory information from different parts of the body, indeed being elicited maximally by the stimulation of a single body part. Consistent with the subjective results, there was neither any significant facilitatory effect of increasing the number of stimulated body parts on threat-evoked skin conductance responses (μS), nor full-body illusion onset times (seconds). In sum, increasing the number of stimulated body parts does not potentiate the full-body ownership illusion; ownership for the whole body does not simply reflect a summation of that across its constituent parts.

Based on this we theorise that to perceive our body as a single unitary whole, a cognitive process specific to this percept must be initiated by, but operate in parallel to, those governing the perception of ownership for the stimulated and non-stimulated body part(s). In addition to the findings discussed above, this conclusion may further be supported by the observation that, although approximately 54.5% of the variance in illusory full-body ownership ratings may be explained by the variance in illusory ownership ratings for all body parts (averaging over both stimulated and non-stimulated body parts), body part ownership cannot explain all of the variance in full-body ownership. Moreover, when analysing the contribution of stimulated versus non-stimulated body parts to the illusory percept for the entire artificial body, we observed evidence of a fairly similar contribution (as indicated by shared variance; McFadden’s pseudo-R^2^) of ownership ratings by the non-stimulated body parts (20.2%), as compared to those body parts that directly received synchronous visuotactile stimulation (33.6%). Therefore, another explanatory variable that can account for the remaining variance in full-body ownership must be involved and may likely reflect cognitive processes that are specific to the full-body ownership percept, possibly involving a unique neural mechanism (Petkova et al. 2011; Gentile et al, 2015). An apt explanation comes from the influential Gestalt psychologist, Kurt Koffka (2013; 1935), who famously stated: “It has been said: the whole is more than the sum of its parts. It is more correct to say that the whole is something else than the sum of its parts, because summing up is a meaningless procedure. Whereas, the whole-part relationship is meaningful”. In line with this conjecture in the visual sciences, in own body perception, the summation of body part ownership does not appear to provide an account of the full-body ownership illusion, whilst the relationships between the parts and the whole were found to be meaningful.

We had a second main aim of painting a more detailed picture of the “spread” of subjective ownership across the entire body by directly comparing illusory ownership for stimulated part(s), non-stimulated part(s) and whole. Hoping to further enlighten important relationships between part- and full-body ownership and gain new insights into how the global feeling of ownership might arise, questionnaire items were designed to separately register illusory ownership ratings toward each of the major body segments in isolation (the right arm, the left arm, the trunk, the right leg and the left leg). We found that the distribution of ownership across the body plan was largely asymmetrical during all of the experimental conditions of the full-body ownership illusion. The two main observations were that 1) body segments were perceived with significantly greater ownership when they were being stimulated synchronously versus when they were receiving no stimulation and 2) for non-stimulated body parts, their location relative to the site of stimulation appeared to influence the magnitude of perceived ownership. For example, there was evidence to suggest that the mannequin’s right hemi-body (often stimulated) was perceived with significantly greater illusory ownership than the left hemi-body (never stimulated). Importantly, however, this was observed even when only the trunk, precisely along the body midline, was stimulated. Similarly, the upper body was consistently perceived with significantly greater ownership than was the lower body; however, further inspection revealed that this finding only characterised the stimulated right hemi-body, and not the unstimulated left. In the healthy population, little research has specifically investigated whether there are any significant differences in perceived ownership between a rubber hand illusion induced on the left versus the right hand. Whilst there is evidence to support a greater illusion susceptibility for participants’ left hand, regardless of handedness (Ocklenburg et al., 2011), others have failed to demonstrate any significant differences in illusion strength owning to whether the left or the right hand is stimulated and whether participants’ are also left- or righthand dominant (Smit et al., 2017; but for proprioceptive drift, see Dempsey-Jones & Kritikos, 2019). Rather than experimentally inducing ownership for a fake hand, Kannape et al. (2019) revealed that the propensity for right-handed participants to experience disownership sensations for their own real hand following the application of asynchronous visuotactile stimulation in a mixed reality paradigm (see Gentile, Guterstam, Brozzoli & Ehrsson, 2013) was significantly increased for the left as compared to the right real hand. However, in the context of illusory full-body ownership perception, to our knowledge, there are no investigations of potential lateralisation effects. Crucially, however, as the present experiment contained conditions applying synchronous visuotactile stimulation to the right hemi-body and never to the left, a possible explanation of the asymmetrical findings in illusory ownership is a covert visuotactile attentional bias to the right hemi-body. Since we did not plan the study to examine possible lateralisation effects, we did not record the participants eye movements during the experiments, nor use a fixation point, relying solely on participants’ adherence to the verbal instructions to attend to the entire body and not to fixate on any particular aspect. However, the findings from the 1S condition fit less well with this explanation and indicate that there might exist a genuine lateralisation effect with slightly weaker illusory ownership ratings for the left hemibody. Future experimental studies should examine this issue because we need to learn more about lateralisation in body ownership illusions and clarify how this relates to literature on lateralisation of body awareness (Tamé et al., 2019; Badde et al., 2019), as well as neurological disorders of body representation (Boll, 1974; Caggiano & Jehkonen. 2018).

In addition to exploring whether increasing the number of body parts receiving synchronous multisensory stimulation would increase the magnitude of the full-body ownership illusion, we also investigated if converging this stimulation across multiple body segments simultaneously could speed up the onset of a full-body ownership percept. Contrary to this, our analyses provided no convincing evidence of significant enhancements to the onset of the full-body ownership illusion owing to whether one, two or three body segments were stimulated synchronously, simultaneously. This finding suggests that there was a stable temporal onset of an illusory full-body percept for all cases of synchronous stimulation, irrespective of the number of receiving body parts. As far as we are aware, the present study is the first that explicitly measure the illusion onset times using a statement that was specifically designed to capture full-body ownership beyond ownership of parts. Previous studies have used wordings such as “please indicate when it feels like the mannequin (or avatar) is your body” and concluded that the onset is rather fast, in the first 10-12 seconds or so (Petkova et al 2011; Preston et al. 2016). However, these studies did not explicitly emphasise the onset of ownership of the entire full body as in the present study. Our onset estimates for Q8 were longer, averaging at 28 seconds for 1S, 25 seconds for 2S and 30 seconds for 3S. Therefore, it seems likely that the participants used a more conservative decision criteria in the present study, only indicating when they indeed experienced ownership for each and every one of the body parts. In earlier studies, participants might have used a more relaxed decision criterion. Compared to the rubber hand illusion, the present onset times for full-body ownership are longer than some estimates of the classic rubber hand illusion (approximately 10s; Ehrsson et al, 2004; Lloyd 2007), but comparable to the average illusion onset time reported for the rubber hand illusion elicited by finger movements; 23 seconds (Kalckert & Ehrsson, 2017). Future studies may endeavour to directly compare ownership for different body parts (e.g. the trunk) and ownership for the whole to gain a better understanding of the temporal relationship between part and full-body ownership. In this way, we may be better able to answer the question: does ownership of parts lead full-body ownership in a systematic way, and are onset times for part and whole correlated? Finally, from a basic method development perspective, we need more data on how good illusion onset times are as a measure of body illusions. For example, how they relate to other measures of the illusion, such as rating scales and objective measures, since it remains unclear whether it is the case that a strong illusion entails a fast onset.

Finally, we explored the relationship between individual variation in Body Awareness Questionnaire (BAQ) scores (Shields, Mallory & Simon, 1989) and susceptibility to the full body ownership illusion. We failed to find evidence of a relationship between this indicator of interoceptive sensibility (Garfinkel et al., 2015) and any of the outcome variables in the full-body ownership illusion. This negative finding seems inconsistent with some studies suggesting a key role for interoceptive processing in the experience of body ownership (Tsakiris et al., 2011; Park & Blanke, 2019; Crucianelli et al., 2013), but in line with other studies presenting results questioning this link (Crucianelli et al., 2018). We reasoned that, as the mannequin does not breathe, the subtle incongruences in felt breathing movements of one’s own chest and the visual impressions of the mannequin’s stationary chest might provide visuo-interoceptive evidence against the full-body ownership illusion. Therefore, we speculated that participants with high BAQ scores could be more sensitive to this type of incongruence. However, our results provided no evidence for such a link. One possibility is that the BAQ may not be sensitive enough to detect the variation in interoceptive sensibility that contributes to the flexibility of full-body ownership during the perceptual illusion. It could also be that interoceptive sensibility itself is less predictive of this variation in general. Other individual differences, for example, those directly related to the multisensory temporal binding window (Shimada et al., 2014; Constantini et al. 2016), may be more fruitful in the future for bodily illusions, as they have been for illusions in the audio-visual domain (Stevenson et al., 2015).

### Limitations of the study

This study focused on the quantification of the subjective experience of part and full-body ownership using questionnaires with rating scales. A limitation of the study was that the objective test of the body illusion, threat-evoked SCR, produced inconclusive results. Although we observed significantly stronger SCR in the 1S condition, the trunk stimulation condition most similar to Petkova and Ehrsson’s (2008) original illusion condition, compared to the 3A condition, the two planned comparisons resulted in more ambiguous results that are difficult to interpret. For example, the 1S condition produced stronger threat-evoked SCR than in the 3S condition, although the questionnaires indicated a similarly strong subjective illusion in these two conditions for both full-body ownership ratings (Q8) and body-part ownership ratings (for the threat-targeted body part, the left leg, Q7). Although we had no reason to think that the 1S could produce the strongest illusion (see introduction), and although there were significant differences between the left and right limbs’ subjective ownership even during 1S, it is possible that objective ownership for the threat-targeted body part, the left leg, was attenuated only in the 3S condition due to the stimulation of the right limbs in 3S and not in 1S. However, more fundamentally, there was no significant difference in threat-evoked SCR between the 3S and 3A conditions, although the questionnaires indicated a very clear and significant difference in both full-body ownership ratings (Q8) and body-part ownership ratings (for the threat-targeted body part, the left leg, Q7). The threat evoked SCR procedure has successfully been used in many full-body illusion studies when contrasting synchronous and asynchronous conditions (Petkova and Ehrsson 2008; Petkova and Ehrsson 2011; Preston et al. 2015; Guterstam et al. 2015; Preuss and Ehrsson 2019) and also in numerous work on the rubber hand illusion (e.g. Armel and Ramachandran; Ehrsson and Petkova 2009; Guterstam et al 2011, 2019). Methodological differences could play a part, as in the present study the SCR data was collected after the questionnaire experiment, which might mean that participants were less alert at this point, and two threat events per conditions were sampled at a fixed time point (at the end of each movie); earlier studies typically included three threat trials per condition presented at unexpected time points and conducted as shorter separate experiments. A more interesting difference is that in the present study the threat was applied to the left leg instead of the abdomen or right arm as in the earlier full-body ownership illusion studies mentioned above. We speculate that the left leg was associated with the smallest ownership-related modulation in threat-evoked SCR between the synchronous and asynchronous conditions, highlighting the need for future SCR experiments that investigate the relationship between part and whole in the full-body illusion from a psychophysiological perspective. Finally, in the SCR experiment, participants were also doing the illusion onset task, and we do not know how conducting that task might have influenced the full-body experience or left leg ownership compared to the initial questionnaire experiments; although, research in the rubber hand illusion suggests that the illusion is not affected by performing cognitive tasks (Fahey et al., 2018). However, it remains that in these previous SCR experiments, the participants did not have any prior task but just relaxed and looked at the body and brushing (Petkova & Ehrsson, 2008). This difference could also be worth pointing out as the participant requirements in the questionnaire experiment and the SCR experiment were slightly different, meaning that it is a little risky to directly compare detailed results across the two experiments.

## Conclusion

The current study supports the feeling of full-body ownership as being mediated by a generalisation of ownership from stimulated part(s)-to-whole, whilst representing an independent percept that may only be subtly enhanced by converging multisensory stimulation across multiple body segments simultaneously. With novel manipulations of the full-body ownership illusion and the future application of modern neuroimaging techniques, we may gain exciting new insights into the neural mechanisms and computations responsible for the experience of full-body ownership in the healthy, adult human brain.

## Additional Information

### Data Availability

Data underlying the study cannot be made publicly available due to ethical concerns regarding participant privacy. This data may be made available pending approval from the ethics committee and would be available upon qualified request to the Brain, Body and Self Lab at.

### Ethical statement

The study was approved by the Swedish Ethical Review Authority.

## Acknowledgements

We would like to thank Martti Mercurio for the Graphic User Interface for the HMDs and his technical support throughout the experiment.

